# Vectorization techniques for efficient agent-based model simulations of tumor growth

**DOI:** 10.1101/032086

**Authors:** Jan Poleszczuk, Heiko Enderling

## Abstract

Multi-scale agent-based models are increasingly used to simulate tumor growth dynamics. Simulating such complex systems is often a great challenge despite large computational power of modern computers and, thus, implementation techniques are becoming as important as the models themselves. Here we show, using a simple agent-based model of tumor growth, how the computational time required for simulation can be decreased by using vectorization techniques. In numerical examples we observed up to 30-fold increases in computation performance when standard approaches were, at least in part, replaced with vectorized routines in MATLAB.

## I. Introduction to the Type of Problem in Cancer

Complex multi-scale cellular automata models are helping cancer researchers to simulate tumor development form the very early stages to clinically apparent disease [1–5]. The main advantages of agent-based models are the ability to formalize single-cell kinetics and to bridge many temporal and spatial scales. However, despite the large computational power of modern computers simulating such complex systems efficiently remains a great challenge. Stochastic simulations often need to be performed hundreds of times in order to estimate parameters during high-dimensional parameter sweeps, or to perform sensitivity analyses of the overall system dynamics. Thus, high-performance implementation techniques are becoming increasingly important. Appropriate use of available data structures and routines can reduce the computational time in some cases more than 80 times [6], making high-dimensional model simulation and analysis feasible.

Agent-based models are commonly implemented in different programming languages including C++, Python or MATLAB. Herein we focus on MATLAB, which is frequently used in scientific computations due to its built-in highly optimized matrix and vector manipulation routines. In particular, we focus on Matlab’s *vectorization* capability, a technique that revises loop-based scalar code into matrix and vector operations. Using a simple agent-based model we show the computational gain when such technique is applied. An initially seeded cancer cell populates the computational domain through migration and proliferation, with spontaneous cell death being possible at mitosis [6].

## II. Illustrative Results of Application of Methods

*In silico* simulations of tumor growth show that the thickness of the proliferating rim and the fraction of quiescent cells depend strongly on the traits of the individual cells. If cells have a short doubling time and small probability to die spontaneously during division, then the proliferating cells are exclusively arranged in a thin rim at the tumor boundary, which amounts to only about 10% of the whole population in our example (Fig. 1A). For slowly dividing cells that are susceptible to spontaneous death during division we observe 2-fold increase in the fraction of proliferating cells in a thicker rim as well as deeper into the tumor (Fig. 1B). Despite the larger proliferative fraction, however, overall tumor growth is slower (45 vs. 69 days to grow from a single cell to a population of 100,000 cells). On the other hand, larger proliferative fraction can result in faster acquisition of favorable phenotypes if spontaneous mutations were considered in the system [4].

**Figure. 1.**
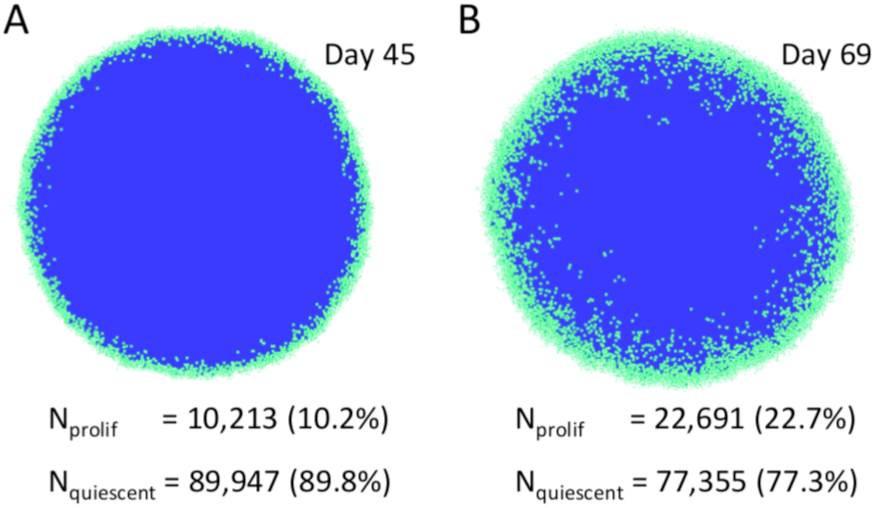
Exemplary tumor morphologies for different single cells traits. (A) Single cell with low doubling time and low chance to die spontaneously during division. (B) Slowly dividing cell with higher chance to undergo spontaneous death during division. Simulation snapshots were taken at the level of about 100,000 cells. Reported are the numbers of proliferating (N_prolif_) and quiescent (N_quiescent_) cells.

We measured the time needed to simulate tumor growth up to about 20,000 cells starting from a single initial cell for three different implementation approaches: 1) loop-based approach, 2) partial vectorization, and 3) extended vectorization. Comparison of the computational speed reveals up to a 30-fold reduction for the extended vectorized approach compared to a loop-based implementation (Fig. 2). Moreover, using appropriate data structures in all three implementation approaches made computational time independent on the lattice size.

**Figure. 2.**
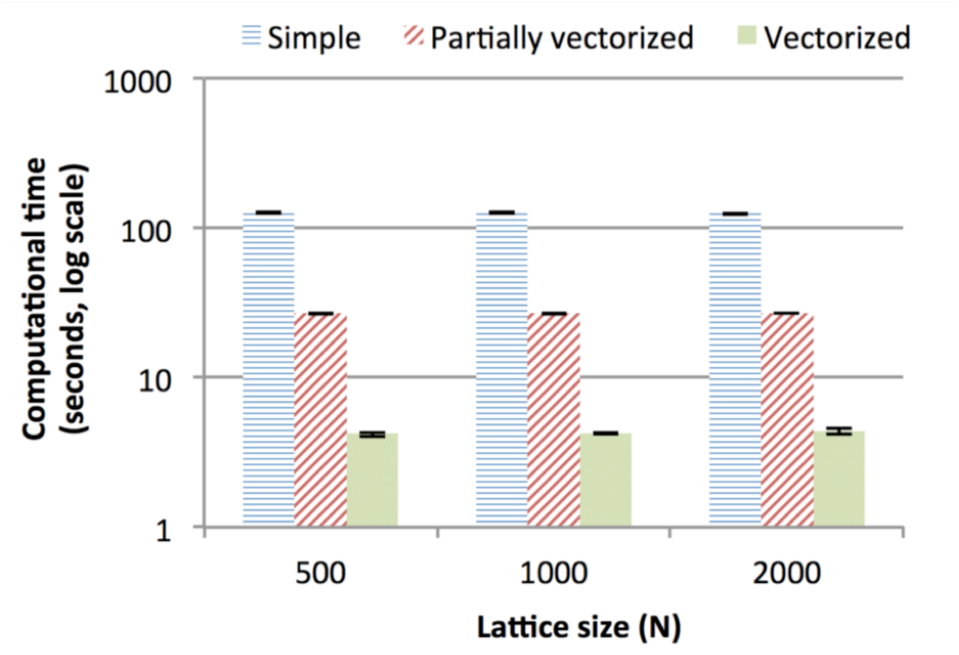
Comparison of the computational speed for different implementation techniques and lattice sizes. Reported are the averages and standard deviations from 20 independent simulations

## III. Quick Guide to the Methods

We consider a simple *in silico* agent-based model, in which each cancer cell occupies a 10×10 μm grid point on a 2D rectangular lattice. Tumor cells populate the computational lattice by cell migration and cell proliferation. A cell can randomly migrate to (with probability p_mig_) or place a daughter cell (with probability p_prol_) into one of the eight adjacent lattice points subject to availability; if all adjacent points are occupied the cell becomes quiescent. Additionally, a cell can die spontaneously at a proliferation attempt (with probability p_death_). Time is advanced at discrete time intervals Δt = 1/24 day (1 hour). At each simulation step, cells are considered in random order to minimize lattice geometry effects and the behavior of each cell is updated.

### A. Basic data structures

We represent the computational domain as an N×N Boolean array (L). A *true* value in L indicates that the lattice point is occupied. An additional integer vector is maintained to store indices for all viable cells present in the system. This additional variable avoids costly lattice searches for existing cells, which otherwise have to be performed at every simulation step. The domain boundaries are set to *true* in L, but not considered in the cells vector.

### B. Basic procedure

A standard loop-based approach to simulate the considered agent-based model is be summarized in the following steps

1. Shuffle cells using the MATLAB randperm function:

~~~
ix = randperm(length(cells));
~~~
2. For each cell check if there is available space in the neighborhood and decide about its fate

~~~
for k = 1:length(ix)
 n = [];
  for i = −1:1
   for j = -N:N:N
    if ~L(cells(ix(k))+i+j)
     n = [n cells(ix(k))+i+j];
    end
   end
  end
 if ~isempty(n)
  %decide about cell fate
 end
end
~~~

Note: variable *n* stores the indices of all free adjacent spots for a currently investigated cell.

### C. Partial vectorization

The nested *for* loops used in the loop-based approach to search through a cell’s neighborhood can be avoided using the following code

~~~
aux = [-N-1 -N -N+1 −1 1 N-1 N N+1];
n = cells(ix(k))+aux(randperm(8));
ind = find(~L(n),1, ‘first’);
n = n(ind);
~~~

Where variable *n* (if non-empty) holds the index to a randomly selected free spot in the cell neighborhood. Of course variable *aux* must be defined outside of the main simulation loop.

### D. Extended vectorization

Rewriting the procedures in a vectorized manner is more involved, but can be summarized in the following steps

1. Select all cells that have a free spot in the neighborhood using vectorized routine

~~~
vnC = 1:length(cells);
aux = [-N-1 -N -N+1 −1 1 N-1 N N+1];
S = L(bsxfun(@plus,cells,aux’));
indxF = vnC(~all(S));
~~~ Variable *indxF* holds the indices of all cells that have at least one vacant lattice spot in their 8-cell neighborhood.
2. Using vectorized logical statements perform initial decisions as to what will happen to each cell

~~~
nC = length(indxF);
P = rand(1, nC)<pprol;
M = ~P & rand(1, nC)<pmig;
~~~ Variables *P* and *M* hold the indices of the cells that will attempt to proliferate and migrate, respectively.
3. Randomly shuffle only those cells that will attempt to proliferate or migrate and, using a *for* loop, determine individual cells fates.

~~~
act = find(P|M);
act = act(randperm(length(act)));
for ii = act
 %decide about cell fate
end
~~~

## Acknowledgment

JP would like to thank Jacob Scott for the inspiration to investigate the advantages of vectorized routines.

